# Taxonomically and metabolically distinct microbial communities with depth and across a hillslope to riparian zone transect

**DOI:** 10.1101/768572

**Authors:** Adi Lavy, Paula B. Matheus Carnevali, Ray Keren, Markus Bill, Jiamin Wan, Tetsu K. Tokunaga, Kenneth H. Williams, Susan S. Hubbard, Jillian F. Banfield

## Abstract

Watersheds are important for supplying fresh water, the quality of which depends on complex interplay involving physical, chemical and biological processes. As water percolates through the soil and underlying weathering rock en route to the river corridor, microorganisms mediate key geochemical transformations, yet the distribution and functional capacities of subsurface microbial communities remain little understood. We have studied metabolic capacities of microbial communities along a meadow to floodplain hillslope transect within the East-River watershed, Colorado, using genome resolved metagenomics and carbon and hydrogen stable isotopes. Very limited strain/species overlap was found at different depths below the ground surface and at different distances along the hillslope, possibly due to restricted hydraulic connectivity after early stages of snowmelt. Functions such as carbon fixation and selenate reduction were prevalent at multiple sites, although the lineages of organisms responsible tend to be location-specific. Based on its abundance, sulfur is significantly more important for microbial metabolism at the floodplain compared to on the hillslope. Nitrification and methylamine oxidation are likely only occurring within the floodplain, with nitrification capacity in shallow soil, and methylamine oxidation in deeper unsaturated sediment. Biogenic methane was detected in deep surface samples, but methanogenic organisms were not identified.

**Originality-Significance Statement:** In a previous study within a hillslope to riparian zone transect of a sub-alpine watershed, the community structure was explored using ribosomal protein S3 genes, and the metabolic potential was hypothesized based on the presence of metabolism related genes. However, tying specific strains and species to metabolic functioning was not discussed as resolved genomes were not available.

In the current study, we use genome-resolved metagenomics along with carbon and hydrogen stable isotopes to explore the spatial distribution of biogeochemical processes. By linking taxonomy and function, using multiple functional genes indicative of full metabolic pathways, we detect heterogeneity in the distribution of metabolic potential and the organisms involved with depth and landscape position. Thus, we infer how microbiome genomic variation impacts biogeochemical cycling across the watershed.

We found very limited strain/species overlap at different depths below the surface and along the hillslope, possibly due to the restricted site to site hydraulic connectivity, and show that communities are largely distinct in their metabolic capacities. Both proximity to the river and the underlying Mancos shale apparently control species distribution and metabolic potential.

Functions such as carbon fixation and selenate reduction were prevalent at multiple sites, although the lineages of organisms responsible tend to be location-specific. Arsenate detoxification was found to be prevalent in the riparian zone whereas selenate reduction was detected within weathered Mancos shale. We conclude that important ecosystem functions are strongly associated with the riparian zone, some of which may have crucial implications as to water quality and human health.

## Introduction

Watersheds, and specifically those of mountains, are important providers of freshwater for areas downstream (Viviroli *et al.*, 2007). These environments are composed of a complex array of ecological life zones and geomorphic units, such as forested alpine and montane meadows and floodplains. These are present in a multitude of geomorphic units, such as mountain tops, hillslopes, and the riparian corridor, and each unit, in turn, has specific subsurface compartments of varying saturation, including soil, weathering zone, and bedrock. Within each geomorphic unit, a multitude of biotic and abiotic interactions occur. Water runoff and infiltration transform and transfer solutes and particles from headwaters to streams and lakes, and from the surface to the subsurface. Currently, there is relatively little known about how hydrobiogeochemical functions are partitioned across ecosystem compartments and the extent to which environmental factors select for distinct microbial communities that mediate biogeochemical transformations and impact water quality and other watershed outputs (Hubbard *et al.*, 2018). Ecosystem gradients are also poorly understood from this perspective, for example how functions change along elevation gradients, with depth below the surface and soil properties. Some prior work examined microbial communities in soils on mountain slopes focusing on the composition of these communities across different climate zones along a transect hundreds of meters long using phospholipid fatty acids (PLFA) detection, or 16S rRNA gene amplification (Djukic *et al.*, 2010; Zhang *et al.*, 2013; Xu *et al.*, 2014; Klimek *et al.*, 2015; Bardelli *et al.*, 2017). While these studies are important for our understanding of climate effects on microbial ecology, they do not address differences at the spatial resolution of meters and centimeters and do not provide metabolic insight for the majority of soil microbes that currently cannot be cultured.

Most studies focus on shallow soil, neglecting deeper soils and weathering rock in mountain ecosystems. The gradients affecting microbial communities within these deeper zones are influenced by above-ground factors such as precipitation, but are also determined by the geochemistry of the underlying bedrock and groundwater (Tytgat *et al.*, 2016; Rempfert *et al.*, 2017). Here, we investigated a well studied headwaters mountainous catchment in the East River of Colorado, where intersecting research activities are focused on developing a predictive understanding of geological, hydrobiogeochemical and ecological process interactions (Hubbard *et al.*, 2018). We analyzed genome-resolved metagenomes to evaluate the connections between microbial composition, metabolic capacities and the spatial organization of these attributes along a hillslope to riparian zone transect and as a function of depth below ground surface.

Given the vast size of watersheds, one way to address the complexity of ecosystem processes is to use a scale adaptive approach in which repetitive elements, or subsystems, are studied separately (Levin, 1992). The scale at which each study is conducted is determined by the hypothesis of the study as well as the methods available for answering it. Following this approach, a “representative” element of hillslope to riparian zone transect was chosen within the Upper East River watershed. From the perspective of vegetation, slope, underlying bedrock and weather conditions, the transect is typical of a substantial fraction of the watershed. The East River watershed in the Upper Colorado River Basin serves as a testing ground for the Department of Energy Watershed Function Project (watershed.lbl.gov) and its associated research community (Hubbard *et al.*, 2018). The watershed is approximately 300 km^2^ with an average elevation of 3266 m. It has an average annual temperature of ~0°C, with average minimum and maximum temperatures of −9.2°C and 9.8°C, respectively. The bulk of the 600 mm yr^−1^ of average precipitation is in the form of snow, which covers the surface between September and May (Pribulick *et al.*, 2016; Hubbard *et al.*, 2018). In early spring, groundwater reaches the surface, but the lowest level of the water table depends on the location along the hillslope. Before the snow melts, the water table is 3.5 m deep at the top of the hillslope whereas at the floodplain it is only ~0.75 m below the surface (Tokunaga et al., under review). The underlying rock at the study site is Cretaceous Mancos Shale, with carbonate and pyrite contents of roughly 20% and 1%, respectively (Morrison *et al.*, 2012). Selenium, which is present in Mancos Shale soils (Elrashidi, 2018), is found at high concentration of 8 ppb in pore waters of unsaturated sediment at the riparian zone whereas sulfate is enriched within groundwater (800 mg/L) at the same location (Lavy *et al.*, 2019).

Microbial communities in soils and sediments play key roles in shaping their surroundings, from rock weathering to nitrogen and carbon fixation (Viles, 2012; Banfield and Nealson, 2018, 12). Prior to the current study, we surveyed microbial metabolic genes from metagenomes that were analyzed without assignment to specific genomes to test the possibility that metabolism of microbial communities varied along the hillslope to floodplain transect. The analysis uncovered indications of trends related to carbon and nitrogen fixation and selenium metabolism (Lavy *et al.*, 2019). These findings motivated the current study, in which the genomes from sample series were comprehensively resolved so that organisms could be confidently linked to functions and the presence of multiple genes required for some functionalities could be verified. Our results reveal that patterns of organism distribution and metabolic potential are shaped by both soil depth and position relative to the riparian zone, and that certain functions are only carried out by distinct organisms at specific sites. We leveraged comparative analyses that depended on the availability of genomes rather than genes to evaluate overlap of strains across sites, addressing the extent to which landscape position contributes to overall watershed biodiversity and constrains inter-site organism dispersal.

## Results

Overall, 6.5 million scaffolds longer than 1 Kbp were assembled from 41 short read datasets from samples collected from between 5 cm and 200 cm depth and distances of 50 - 280 m from the river (Table S1). Given that microbial communities in soils are diverse and therefore difficult to resolve genomically, it is unsurprising that, on average, only 27.8% (±11) of the reads could be mapped back to these scaffolds. The large fraction of unmapped reads reflects the huge microbial diversity in soil and the prevalence of organisms at similar very low abundance. Scaffolds were binned into draft genomes, 1448 of which were assessed to be ≥60% complete based on the presence of 43 bacterial single copy genes or 38 archaeal single copy genes. The bins contain 4.05 Gbp of DNA sequence in 625,348 scaffolds (~10% of all scaffolds). Dereplication at 98% average nucleotide identity (ANI) reduced the number of genomes to 1005, and screening for genomes that are at least 80% complete resulted in 484 unique near-complete genomes (Table S2). These genomes were classified as high-quality draft (111 genomes) or medium-quality draft (350 genomes) according to the completeness and redundancy scores used by Anvi’o (Eren *et al.*, 2015), and the genome reporting standard (Bowers *et al.*, 2017). These values are reasonably high, considering the strain complexity of microbial communities in soil, which results in genome fragmentation (Myrold *et al.*, 2014). Twenty-three draft genomes had redundancy score higher than 10 and therefore did not match any of the categories in the reporting standard. It should be noted that Anvi’o generates a redundancy value which does not completely correlate with the definition of ‘contamination’ as used in the genome reporting standard.

Genomes of organisms from 28 phyla and 5 proteobacterial classes were assembled from the hillslope to riparian zone transect. Among the genomes were those of organisms typically detected in soils in 16S rRNA gene surveys (Janssen, 2006). These include *Acidobacteria*, *Actinobacteria*, *Bacteroidetes*, *Planctomycetes*, *Proteobacteria*, and *Verrucomicrobia* (Figure S1). Genomes representing *Woesebacteria*, *Yanofskyabcteria*, and *Wolfebacteria* from the Candidate Phyla Radiation group were reconstructed. These were previously detected only at the floodplain by their rpS3 gene, and most often in samples from 10 cm above the water table (Lavy *et al.*, 2019). Having genomes of taxa which were previously only detected by a marker gene presents an opportunity to explore their potential metabolism and environmental impacts.

Two clades of *Deltaproteobacteria* displayed a differential preference for either the floodplain or the hillslope. The first clade, present at the floodplain, consists of *Desulfuromonadales* (*Geobacteraceae*), Syntrophobacterales (*Syntrophaceae* and *Syntrophobacteraceae*), *Thermodesulfobacteriales* (*Thermodesulfobacteriaceae*), and unknown *Deltaproteobacteria* mostly similar to those genomically described from Rifle, CO. These species were found to be significantly correlated with water-saturated sediment and from 10 cm above the water table. The second clade, containing species of Myxococcales (*Cystobacterineae*, *Nannocystineae*, and *Soranglineae*), was present at all parts of the transect other than water-saturated sediment. Actinobacteria, which are routinely found in soils and represented here by 70 genomes, were significantly correlated only with hillslope sites, although some were detected at the riparian zone.

The presence of strain level, nearly identical genomes across the transect requires relatively recent inter-site transport, probably via downslope subsurface water flow. Conversely, lack of strain sharing could be explained either by limited inter-site dispersal or selection against genotypes adapted to conditions elsewhere on the hillslope. Only seven bacterial and two archaeal strains occurred at more than one sampling location (Figure 1). One strain of *Actinobacteria* (Ac2), was found at two sites that are not adjacent (PLM1 and PLM6), but at a similar depth, 60 and 50 cm, respectively.

**Figure 1.**
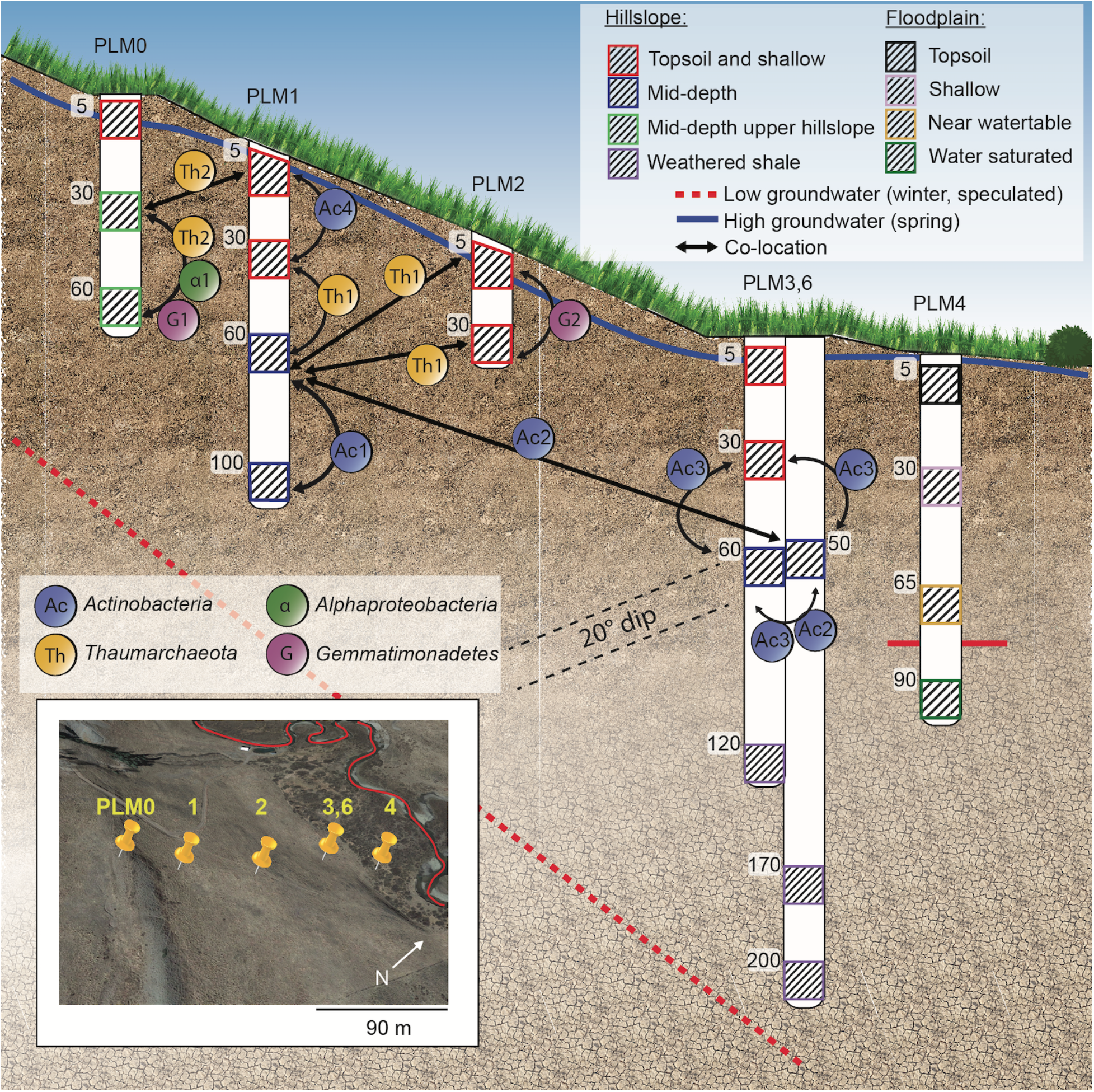
Sampling scheme, connectivity between sites and samples grouping. Numbers along wells indicate depth below-ground surface of sample (in cm). Specific depths that were sampled within each PLM site along the transect are marked with diagonal lines pattern. Nine strains were found to occur at more than one sample. These are from the class *Alphaproteobacteria*, and *Thaumarchaeota*, *Actinobacteria*, and *Gemmatimonadetes* phyla. Identical strains were detected within adjacent depths of a site, or between two adjacent sites. Only Ac2 co-occurred in two sites that are not adjacent (PLM1 and PLM6) but at similar depths, 60 and 50 cm, respectively. Samples similar in community composition, as was described by Lavy et al. (Lavy *et al.*, 2019), were grouped to form a single zone which is denoted by a colored box. A 20° dip towards the South-East is depicted by dashed lines. An aerial photo of the transect is shown within the inset.

Drilling and installation of sampling equipment for another experiment at each of the PLM locations revealed compositionally distinct shale layers with a 20° dip to the southwest, paralleling the prominent lithological feature expressed at the surface at PLM0. Thus, lithological variation in the Mancos shale should be considered when evaluating the effect of depth below the surface on community composition along the hillslope transect.

We searched the 484 representative genomes for 23 metabolic functions associated with five types of reactions, namely C1 metabolism, oxidation processes (electron donors), reduction processes (electron acceptors), carbon and nitrogen fixation, and resistance to toxic elements. The genomes were found to harbor 20 out of the 23 metabolic functions. Denitrification, methane oxidation, and respiratory arsenite oxidation were absent from the draft genomes as they lack the required genes that encode for enzymes involved in these pathways. Among all electron acceptors, organisms potentially capable of using oxygen for respiration were the most abundant. However, genomes of organisms that could potentially respire selenate and sulfite had high relative abundance within water-saturated sediment at the floodplain. Organisms with the potential for sulfur respiration were not detected at any of the hillslope sections (Figure 2).

**Figure 2.**
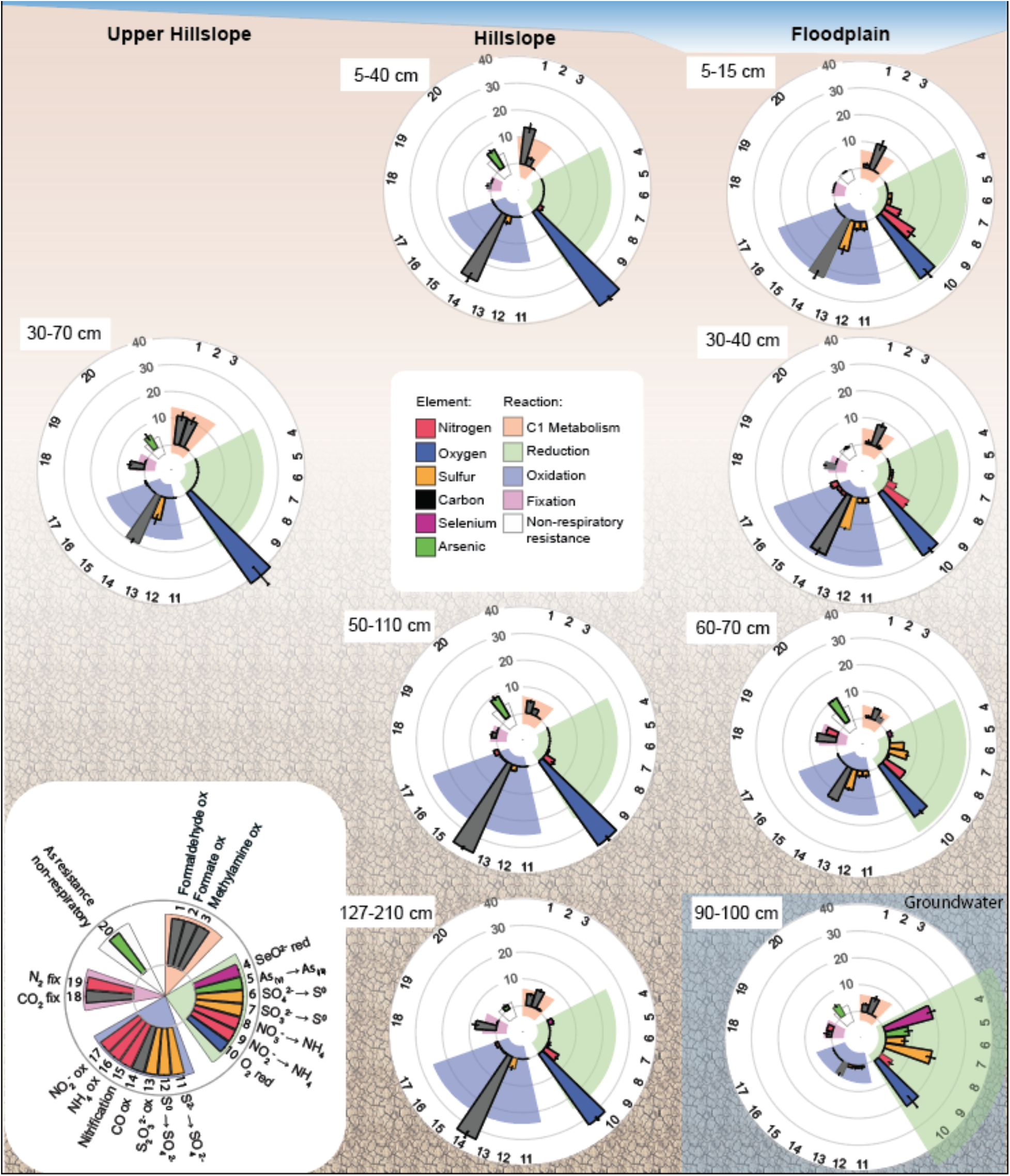
Relative abundance of metabolic functions at each zone. The metabolic potential of each genome was evaluated using Hidden Markov Models for key enzymes of metabolic function. The color of each bar represents the element which is directly involved in the metabolic pathway. The type of reaction (i.e., C1 metabolism, reduction, oxidation, fixation or non-respiratory resistance) is denoted by the color of background behind bars. The length of the reaction type bar (e.g. C1 metabolism) is correlated with the overall relative abundance of functions within the category and is scaled to a maximum of 60. Bars represent SE. Legend is given at the lower left corner of the figure.

By using genomes rather than individual genes without genomic context, we were able to show that the relative abundance of organisms capable of using sulfate and sulfite as terminal electron acceptors increases with depth at the floodplain, whereas the relative abundance of those capable of oxidizing thiosulfate, sulfur and sulfide diminish (Figure 2). This follows the general trend wherein the relative abundance of organisms with reductive functions increases with depth where the trend for oxidative functions decreases. Non-respiratory arsenic resistance, either via redox or methylation or arsenic species is the third most abundant function at 10 cm above groundwater. However, its relative abundance is much lower just 20 cm below the water table, and within weathering shale at the hillslope. Interestingly, we detected organisms capable of dissimilatory arsenate reduction using the *arr* operon, but could not detect the full sets of genes which make the *arx* operon, encoding for anoxygenic arsenic oxidation. Deltaproteobacteria that are relatively abundant in the riparian zone sediments encode an arsenate reductase *arr* operon (a third partial *arr* operon, including the key *arrA* subunits and *arrS* was also detected). The highest abundance of genes encoding for arsenite methylation detoxification that may follow arsenate reduction were detected within unsaturated sediments at the floodplain (Figure S2). These genes were also found in every compartment other than shallow soil of the floodplain.

Applying the indicator species analysis to metabolic functions allows identifying functions that uniquely describe one or multiple sites. Organisms with nitrogen fixation genes were most abundant in samples taken from near groundwater, above and below the water table, but not in other samples. Therefore, nitrogen fixation was found to be significantly representative of these zones (Indval = 0.983, p-value = 0.002) (Figure 3). Organisms capable of oxidizing ammonia to nitrite were abundant at several zones but the process was identified as an indicator process only for floodplain-associated shallow samples (Indval = 0.764, p-value = 0.033). Sulfite and sulfur reduction as well as sulfur and sulfide oxidation were indicators of any sample at the floodplain as these functions were absent from the hillslope. While organisms with the potential to fix CO_2_ were detected in every sample other than the topsoil of the floodplain, these were most abundant in weathered shale and in the unsaturated zone near the water table.

**Figure 3.**
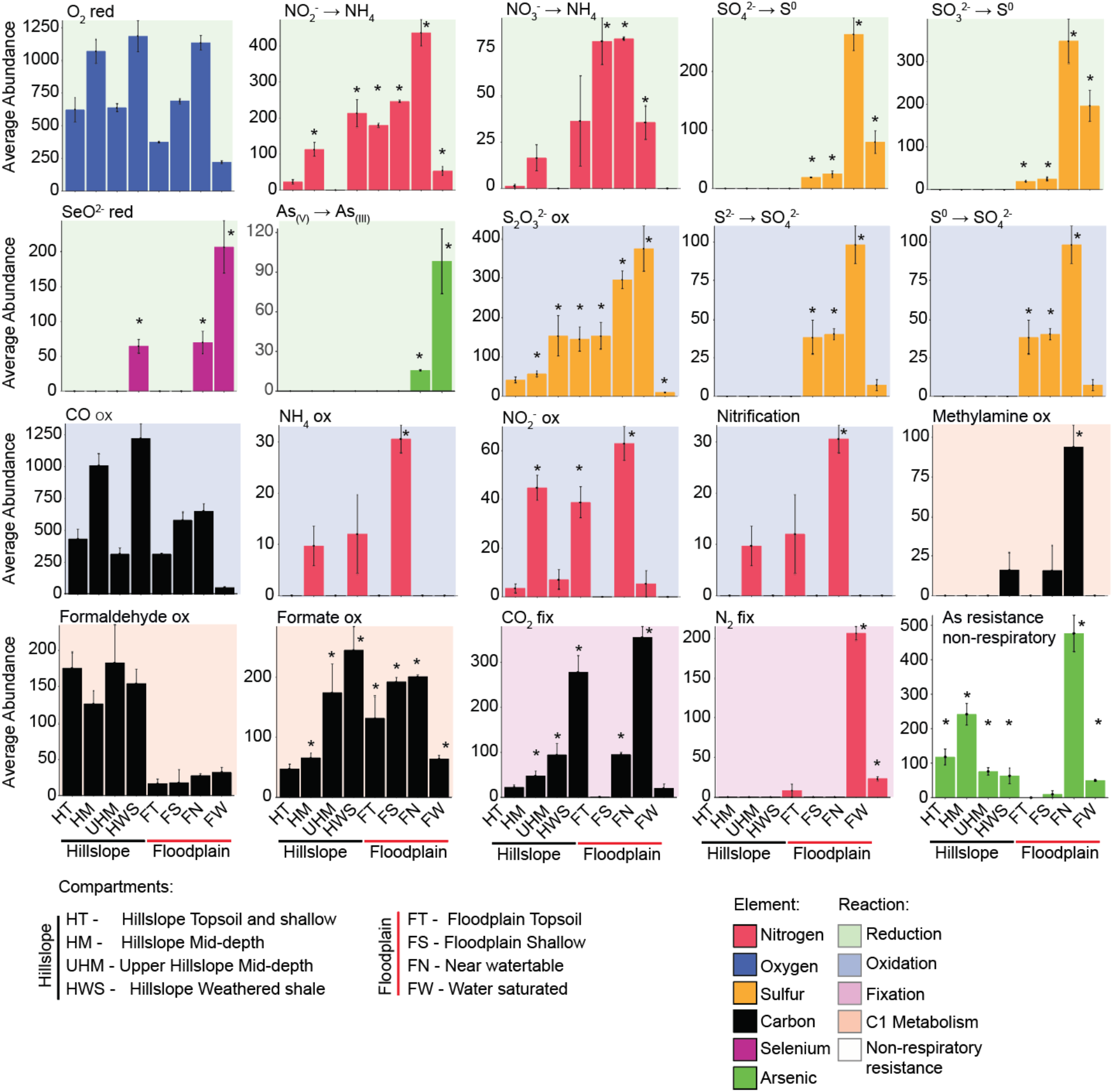
The abundance of functions and indicator functions. Bars marked with stars are indicator functions for the samples. For example, nitrification is an indicator function at the near water table zone, whereas carbon fixation is an indicator function for at multiple zones (hillslope mid-depth, upper hillslope mid-depth, weathered shale, floodplain shallow samples, and floodplain near water table). The color of each bar represents the element which is directly involved in the metabolic pathway. The type of reaction (i.e., C1 metabolism, reduction, oxidation, fixation or non-respiratory resistance) is denoted by the color of background behind bars. Bars are SE. Representation of each sample is given in Figure S4.

Organisms predicted to oxidize formaldehyde were more abundant at the hillslope compared to the floodplain. At hillslope sites, formaldehyde oxidation independent of thiol and dependent on tetrahydromethanopterin (H4MPT) and glutathione or Bacillithiol/Mycothiol appears prominent (Figure S3). However, a fused gene encoding for a bi-funcitonal enzyme with formaldehyde activating enzyme (fae) and hexulose-6-phosphate synthase (hps) capacity was found within the genomes of *Bathyarchaeota*, which were detected in groundwater saturated sediment at the floodplain.

Methanogens were not detected in the microbial communities at the studied sites. However, concentrations greater than ~3200 mg/L_H2O_ of methane were measured in the water phase at the PLM6 well from January to July 2017, and concentrations greater than ~ 1100 ppm_v_ were observed in the gas phase in an adjacent well (PLM3) in May and July 2017 (Table S3). Carbon and hydrogen stable isotopes of methane show mixing between acetoclastic and hydrogenotrophic methane, except for samples taken from 3.7 meters at PLM3 where methane has isotopic ratios typical of hydrogenotrophic methane (Figure 4).

**Figure 4.**
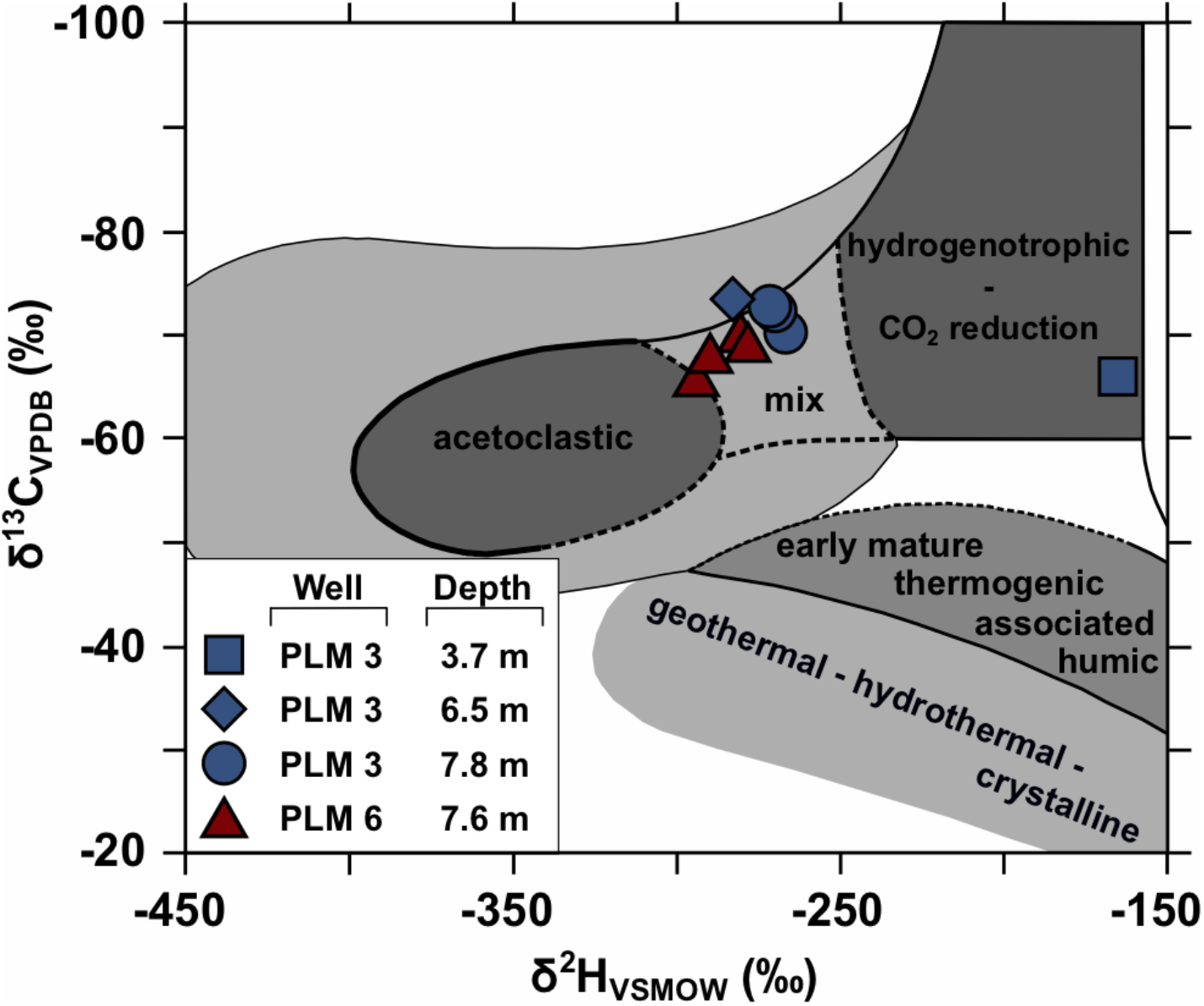
Carbon and hydrogen isotopic compositions of methane samples. δ^13^C and δ^2^H values indicate mixing between acetoclastic and hydrogenotrophic methane for depths from 6.5 m to 7.8 m. Well PLM3 at 3.7 m shows hydrogenotrophic methane. This figure was adapted from Whiticar (1999).

Some microbially mediated biogeochemical reactions are predicted to occur at multiple study site locations. However, the processes are apparently mediated by organisms from different taxonomic lineages. For example, *Deltaproteobacteria* (*Desulfobacca acetoxidans*, *Syntrophorhabdus aromaticivorans*, *Geobacteraceae*, and species closest to *Deltaproteobacteria* RBG genomes) are potential reducers of selenate at the hillslope, whereas Candidatus *Methylomirabilis oxyfera*, and *Rokubacteria* conduct the same role at the floodplain (Figure 5). The selenate reductase gene *srdA* in *Rokubacteria* was found on a 4.05 Kbp scaffold also carrying genes encoding a putative polysulphide reductase *NrfD* family protein, and 4Fe-4S ferredoxin-type, iron-sulfur binding domain, both of which are likely components of the microbial electron transport chain.

**Figure 5.**
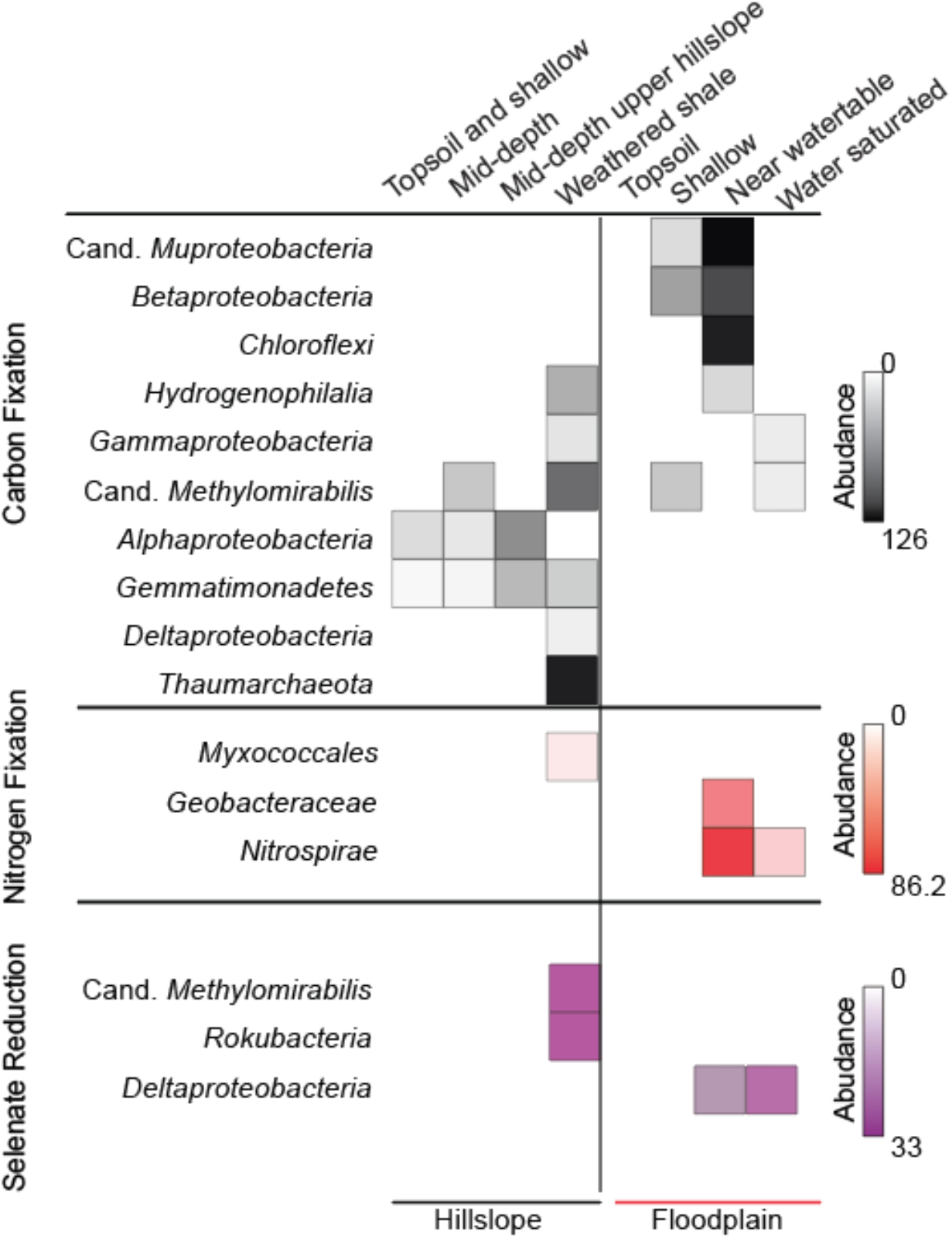
Metabolic functions conducted by different taxa at the hillslope compared to the floodplain. CO_2_ fixation, N_2_ fixation, and selenate reduction all occur both at the hillslope and floodplain zones. However, these functions are carried out by organisms from different taxa at each location. Heatmap shows the average abundance of organisms with the potential of these functions at each zone.

Patterns of partitioning for N_2_ and CO_2_ fixation are similar to those for selenate reduction. All genes required for nitrogen fixation are present in the genomes of *Myxococcales* (class *Deltaproteobacteria*) in the weathered shale zone from the hillslope, whereas at the floodplain this role is taken by *Geobacteraceae* and *Nitrospirae* closely related to species from Rifle, CO (Anantharaman *et al.*, 2016). The abundance of organisms that may fix CO_2_ increases with depth at the hillslope as well as in the floodplain, except for within groundwater saturated samples. The most abundant autotrophs occur exclusively at the hillslope and are *Alphaproteobacteria, Gemmatimonadetes*, *Deltaproteobacteria, and Thaumarchaeota*. While *Thaumarchaeota* species seem to fix CO_2_ via the 3-hydroxypropionate /4-hydroxybutyrate (3HP-4HB) pathway, species of the other phyla may use the non-photosynthetic Calvin cycle for this purpose. At the floodplain, *Betaproteobacteria*, *Chloroflexi*, and Candidate phyla *Muproteobacteria* may conduct carbon fixation utilizing the non-photosynthetic Calvin cycle.

## Discussion

In a previous study conducted at the same site, and in which only rpS3 protein sequences were used as taxonomic markers, organisms from 37 phyla were detected (Lavy *et al.*, 2019), although, candidatus Lambdaproteobacteria and candidatus Muproteobacteria are arguably part of the Proteobacteria (super)phylum. In the current study we reconstructed genomes for bacteria from 25 phylum-level groups. The phylum-level lineages for which genomes were not recovered are *Amesbacteria*, *Katanobacteria*, *Latesbacteria*, *Modulibacteria*, *Moranbacteria*, *Nealsonbacteria*, *Parcubacteria*, and *Yanofskybacteria* of the Candidate Phyla Radiation CPR) and *Tectomicrobia*. We attribute the lack of genome recovery to the low relative abundance of these bacteria (<0.04%, Lavy *et al.*, 2019). Additionally, we did not recover genomes of *Spirochaetes,* which were detected in 8 out of 41 samples and had a maximum relative abundance of ~1% based on the rpS3 gene (Lavy *et al.*, 2019), or for members of the Candidate phylum *Lambdaproteobacteria*. The detection of organisms for which genomes were not recovered is unsurprising due to the challenges of resolving genomes from complex communities such as found in soils (Myrold *et al.*, 2014).

The sampling sites along the transect follow a hydraulic flow line that connects sites from a higher elevation with their downstream counterparts (Tokunaga *et al.*, 2019). The sites are occasionally connected by infiltration and lateral subsurface flow but rarely by overland runoff. During this late summer investigation, before snowfall had started and when vertical and lateral waterflow did not occur, the availability of genomes allowed us to establish that microbial communities at different sites share very few organisms. Additionally, while soil and saprolite at the site are derived from Mancos shale (Morrison *et al.*, 2012), local differences in the physical and chemical properties of the parent rock could impose different selective pressures on microbes. This could contribute to taxonomic variability at similar depths but at different sites. The finding of many genomes from the same species (but not strains, see Figure 1) in multiple sites suggests that similar, but not identical, environmental conditions select for similar, but distinct organisms. However, the observation that 8 of 484 genomes are from near-identical strains that occur at two depths or at more than one site, probably requires dispersal through the soil and saprolite profile. The finding of genotypically near-identical *Actinobacteria* and two *Thaumarchaeota* in sites downslope from each other probably indicates dispersal between sites across the hillslope. If transfer occurs, it is probably during the weeks following snowmelt, when groundwater flows, but at only ~10 to 20 m per month (Tokunaga *et al.*, 2019)

Beyond strain tracking, the strength of the genome-resolved metagenomic approach lies in the potential for metabolic inference. Metabolic potential is not confounded by temporal fluctuations in physical and chemical conditions that impact transcript-based assays. Although many scaffolds were unbinned, the availability of genomes for most of the most abundant organisms in the environment suggests the possibility for metabolic prediction for those most likely to most influence overall biogeochemistry at the sites from which the genomes were recovered.

Microbes and hydrology were suggested to jointly control the fate of carbon and nitrogen in the East River watershed (Newcomer *et al.*, 2018). Organisms capable of CO_2_ fixation were detected throughout the hillslope, at highest abundance within weathered shale (127-200 cm depth) and linked to *Thaumarchaeota*, and at every depth other than the shallow zone at the floodplain, indicating potentially important microbial contributions to soil and sediment organic carbon compound inventories. While organic carbon from decomposition of organic material is readily available for microorganisms in shallow soil, CO_2_ fixation could sustain heterotrophs in the subsurface. Members of *Betaproteobacteria* and *Chloroflexi* were previously described to be autotrophs (Howarth *et al.*, 1988; Hou *et al.*, 2018), but carbon fixation by Candidatus *Muproteobacteria* is suggested here for the first time. Environmental conditions probably select for different bacteria/archaea with these roles under different conditions.

The presence of methylamine oxidizing organisms is significantly correlated with unsaturated sediment from above the water table at the floodplain, but this capacity was also detected in shallow soil at the floodplain, but at lower abundance. Methylamine is released from biodegradation of proteins and nitrogen-containing osmolytes (Barrett and Kwan, 1985), for example, those released from microbial cells due to repeated drying and rewetting (Halverson *et al.*, 2000), and can be used as sole carbon and energy source for methylotrophs (Anthony, 1982), as well as a nitrogen source (Taubert *et al.*, 2017). The origin of methylamine in the deeper zones of the floodplain is unclear, but it could be sourced from the surface or formed by oxidation of compatible solutes and other compounds from microbial community members (Daly *et al.*, 2016).

Biogenic methane was detected in hillslope groundwater at depths greater than those that were sampled for metagenomic analysis. The finding that methane from 3.7 m depth was hydrogenotrophic, but of mixed biogenic origin at other sites, is interesting because hydrogenotrophic methanogenesis, in which H_2_ (or sometimes formate) is used as an electron donor (Liu and Whitman, 2008; Thauer *et al.*, 2008), is the most widespread methanogenic pathway (Thauer *et al.*, 2008). These archaea reduce CO_2_ to CH_4_ by utilizing the reductive acetyl-CoA or Wood–Ljungdahl pathway (Berghuis *et al.*, 2019). In order to conserve energy, hydrogenotrophs may couple the Wood–Ljungdahl pathway to methanogenesis. While genes responsible for formate oxidation were detected in some genomes, these organisms did not have the required genes for a complete methanogenesis. Therefore it is suggested that methanogenic archaea are predominantly confined to the groundwater at greater depths than studied here. Similarly, the capacity for methane oxidation could not be established. However, 24 genomes encode *amoABC/pmmoABC* genes, but structural similarities between the ammonia oxidizing and the particulate methane oxidizing variants, preclude assignment to a specific function.

Some organisms have genes required to oxidize formaldehyde and formate, which may be formed by methane oxidation or by fermentation and breakdown of plant and animal products (White, 2007) such as methanol. Formaldehyde oxidation could be a means of detoxification (Thauer *et al.*, 2008), but could also be used for energy production and as a carbon source via the ribulose monophosphate (RuMP) pathway (Thauer *et al.*, 2008). The presence of a fused gene encoding for a bifunctional enzyme with formaldehyde activating enzyme and hexulose-6-phosphate synthase exclusively in organisms from groundwater saturated sediment may indicate that formaldehyde is an important carbon source within this ecological compartment.

Nitrogen fixation often takes place in estuarine, coastal environments, and river banks sediment (Howarth *et al.*, 1988). Specifically, at river banks, nitrogen fixation could account for a significant portion of nitrogen transported to groundwater. For example, 9.3% of the total terrigenous inorganic nitrogen transported into the Yangtze estuarine and coastal environment is attributed to biological (as opposed to industrial) nitrogen fixation (Hou *et al.*, 2018). At the East River study site, nitrogen fixation at the floodplain, particularly close to groundwater, could contribute to the flux of nitrogen into the river, affecting downstream water quality. The identity of nitrogen fixers is differentiated spatially, probably due to selection for other traits. The reducing environment in water-saturated sediments may explain why nitrate-reducing, and to a larger extent nitrite reducing organisms were more abundant at the floodplain compared to the more aerobic environment at the hillslope. Some degree of stratification involving nitrogen compound reduction capacities in shallower regions and sulfate reduction in deeper regions of the profile that spans aerobic, transitional and groundwater saturated regions of riparian zone extends similar findings related to stratification in marine or terrestrial sediments (Smith and Harris, Jr., 2007; Wright *et al.*, 2012).

A previous study at the East River watershed suggested that the loss of sulfate across a low-gradient, meandering section of the watershed could indicate microbial sulfate reduction in floodplain sediments (Carroll *et al.*, 2018), and this process was suggested to modify the hillslope inputs of sulfur species to the riparian zone. The abundance of organisms in the water-saturated zone predicted to be capable of reducing sulfate and sulfite to elemental sulfur, supports this suggestion.

Arsenic is present in the Mancos shale, primarily as an impurity in pyrite (FeS2) (and selenide) minerals, and is released to solution as the trivalent form, arsenite, or the pentavalent form, arsenate, following oxidative dissolution of pyrite during shale weathering (Plant, J.A. *et al.*, 2003). Microorganisms can alter arsenic speciation and transform arsenic compounds between biotic (i.e., methyl-arsenic and arsenosugars) and abiotic forms (Oremland and Stolz, 2005). Arsenate may be reduced to the more toxic form, arsenite (Oremland and Stolz, 2005) with or without methylation (Zhu *et al.*, 2017) prior to export from cells during detoxification.. Two genomes of Deltaproteobacteria that are relatively abundant in the riparian zone sediments encode an arsenate reductase operon. Thus, these bacteria may decrease arsenate and increase arsenite concentrations in groundwater and the river. In unsaturated sediment near the water table, genes involved in arsenite methylation were detected at high abundance, suggesting that arsenic might be methylated before being released into the river. These processes could impact downstream water quality.

Selenium (Se), an essential element for several enzymatic reactions, becomes toxic when present at a high concentration in its most oxidized form, selenate. Se occurs in Mancos shale in the form of insoluble metal selenides (Elrashidi, 2018) that can be oxidized, forming mobile selenite and selenate (Presser, 1994; Gebreeyessus and Zewge, 2018). Here, we attribute reduction of selenate released by weathering of the Mancos shale to *Deltaproteobacteria* within the floodplain and to *Rokubacteria* and Candidatus *Methylomirabilis* at hillslope sites. *Geobacter* species (phylum *Deltaproteobacteria*) are known to reduce selenite (Pearce *et al.*, 2009) and Candidatus *Methylomirabilis* may be capable of coupling anaerobic methane oxidation to selenate reduction (Luo *et al.*, 2018). However, this is the first evidence suggesting that *Rokubacteria* is capable of selenate reduction. The activities of these three groups of bacteria have the potential to limit the export of toxic, mobile selenate from the watershed.

Riparian ecosystems provide key services to society (González *et al.*, 2017). Some studies suggest that they are particularly sensitive to climate change impacts (Tockner and Stanford, 2002; Perry *et al.*, 2012; Capon *et al.*, 2013) whereas other studies predict that riparian zones will be relatively resilient, as they evolved under conditions of high environmental variability (Catford *et al.*, 2013). Within mountainous watersheds such as studied here, rising temperatures and atmospheric CO_2_ concentration, droughts and early snowmelt all have the potential to alter the riparian ecosystem (Hubbard *et al.*, 2018). This may alter water table elevations, and consequently depth of sub-surface anoxia, affecting vegetation (Vartapetian and Jackson, 1997) as well as microbial community composition and functioning (Rühle *et al.*, 2015). Lowering of the water table could accelerate the release rate of solutes, such as nitrate, sulfate, iron, and selenate from soils, causing a deterioration of water quality both locally and downstream (Freeman *et al.*, 1993). The chemical forms of these elements, as well as the reaction rates, are, in part, determined by transformations mediated by subsurface microbial organisms. The distribution of functions of different organisms (and in different metabolic contexts) could provide functional flexibility as conditions change over the course of the year and support system resilience to longer-term environmental changes (Allison and Martiny, 2008). Our findings of functional differentiation between the hillslope and the riparian zone and further partitioning across the hillslope constrains understanding of the watershed ecosystem.

## Conclusion

Most elemental cycles near the Earth’s surface are microbially mediated, but we know very little about how these biogeochemical processes are distributed across organisms and environmental compartments, especially at the watershed scale. Here, we show that such questions can be addressed across large transects, from high on hillslopes down to the river corridor, via genomic analysis of microbial communities. A similar methodology could be used across scalable transects in other environments. We find heterogeneity in the distribution of metabolic potential, the organisms involved, and presumably in the biogeochemical cycling that occurs at each site. Thus, we conclude that different ecosystem compartments such as the riparian zone and weathered bedrock need to be identified and studied individually. However, sites must also be studied in combination to begin to predict how nutrients and contaminants released from underlying rock are modulated by spatially variable biological processes to dictate exports from watersheds.

## Experimental Procedures

### Sampling

The Pumphouse Lower Montane (PLM) intensive study site is located on the north-east facing slope of the East River valley near Crested Butte, Colorado, USA (38°55’12.56”N, 106°56’55.39”W). Soil and sediment samples were collected as described in Lavy et al. (Lavy *et al.*, 2019). Briefly, samples were taken during four days in June 2016 from five sites along a hillslope transect and one site at a floodplain using manual augur and sterile plastic liners. These sites ordered from highest to lowest elevation are PLM0, PLM1, PLM2, PLM3, and PLM4. All samples other than those taken at 90 cm depths at PLM4 were from above the water table. An additional site at the floodplain was sampled using an air-cooled split-spoon Odex drill during the same period of time. The installation of sampling equipment for another experiment at the same locations revealed compositionally distinct shale layers over the hillslope with a 20° uplift to the northeast. The first five centimeters of topsoil and plant material were cleared from the surface of each sampling site before samples were taken. In total, 20 samples were collected as follows: PLM0 - 5, 30, 60 cm; PLM1 – 5, 30, 60, 100 cm; PLM2 – 5, 30 cm; PLM3 – 5, 30, 60, 127 cm; PLM6 – 50, 170, 200 cm; PLM4 – 5, 32, 65, 90 cm. Each sample was manually homogenized in a sterile Whirl-Pak bag and aliquots of 5 g cleared from rocks and roots were placed on ice for DNA extraction, as well as in 10 ml of LifeGuard Soil Preservation Solution (Qiagen, Netherlands) for RNA and DNA co-extraction.

### DNA extraction and sequencing

The extraction process is described in Lavy et al. (2019). Briefly, DNA was extracted from 10 g of soil or sediment with DNeasy PowerMax Soil Kit (Qiagen, Netherlands) in two batches of 5 g each which were combined during the cleaning step. DNA was also co-extracted with RNA from 5 g of soil using RNeasy PowerSoil Total RNA Kit (Qiagen, Netherlands) and Phenol:Chloroform:Isoamyl Alcohol 25:24:1 saturated with 10 mM Tris (final pH 8.0) and 1 mM EDTA. RNeasy PowerSoil DNA Elution Kit (Qiagen, Netherlands) was used to collect DNA which was further cleaned using DNeasy PowerClean Pro Clean Up Kit (Qiagen, Netherlands). Overall, two DNA samples were produced from each sampling, one from DNA extraction and the second from the DNA that was co-extracted along with RNA. A third DNA sample was extracted from the 90 cm deep PLM4 sample, thus a total of 41 DNA samples were used for further analysis.

Metagenomic libraries were prepared at the Joint Genome Institute (JGI) after validating concentrations and DNA integrity using Qubit (Thermo Fisher Scientific) and gel electrophoresis, respectively. NEB’s Ultra DNA Library Prep kit (New England Biolabs, MA) was used for library preparation for Illumina with AmpureXP bead selection aimed for an average fragment length of 500 base-pair (bp) following the manufacturer’s protocol. The libraries were sequenced at JGI in an Illumina Hiseq 2500, generating 150 bp paired-end, sequences (Table S1)

### Sequences analysis and genome binning

Reads processing, scaffolds assembly, and open reading frame annotation were done as reported in Lavy et al. (2019). Genomes were binned manually using the ggKbase platform (Jillian F Banfield, 2015) as well as with the automated binners CONCOCT v0.4 (Alneberg *et al.*, 2014), Maxbin2 (Wu *et al.*, 2016), Abawaca1, Abawaca2 (Brown *et al.*, 2015), and MetaBAT v0.32.4 (Kang *et al.*, 2015). All predicted bins from each sample were used as input for DAStool v1.1.0 (Sieber *et al.*, 2018) which selected for the best representative genome. Representative genomes from all samples were then dereplicated with dRep v2.2.2 (Olm *et al.*, 2017) at 98% average nucleotide identity (ANI), which selected for the best species level representatives. Each of the resulting genomes was further curated in ggKbase, and genomes considered as near-complete, having ≥ 80% of 43 bacterial single copy genes or 38 archaeal single copy gene sets (Anantharaman *et al.*, 2016) were selected for further curation. These genes were imported into Anvi’o v5.4 (Eren *et al.*, 2015) for further manual inspection and refinement using differential coverage, kmer frequency, and GC content.

### Genome abundance across samples

To determine the presence and abundance of genomes across samples, reads from each sample were mapped against genomes with bowtie2 (Langmead and Salzberg, 2012). The average coverage and breadth of coverage (percent of genome length with a coverage greater than 1) of each genome in each sample was then calculated (Olm et al., 2017). Each genome is considered to be present in at least one sample (at a minimum, the sample from which it was originally binned) but could be falsely identified in other samples due to a low breadth cutoff (i.e., false positive). Therefore, we implemented a breadth cutoff of 0.828 based on the lowest breadth cutoff that retains all genomes from the samples they were originally binned from (Figure S5). The coverage of the genomes was adjusted according to the number of basepairs that were sequenced from each sample to compensate for differences in sequencing depth.

### Strains detection

Strains of the same species were identified as either genome that shares more than 99% ANI according to their dRep cluster placement (Olm *et al.*, 2017), or by having an average breadth > 0.99 within more than one sample.

### Partitioning into zones

Samples were grouped based on the results of a previous study (Lavy *et al.*, 2019) in which community composition was defined by ribosomal protein S3 (*rpS3*), and in which microbial communities at the hillslope and floodplain compartments were found to be distinct. Furthermore, a depth gradient was observed to affect community composition and potential metabolism. Therefore eight zones were defined for the purpose of the current study. At the hillslope compartment, samples were grouped into either Top and shallow soil (n=14), mid-depth zone (n=8), mid-depth at an upper hillslope location (n=4), or weathered shale (n=6) (Figure 1). At the riparian zone compartment, samples were grouped into either topsoil (n=2), shallow soil (n=2), unsaturated sediment at 10 cm above the water table (n=2), or groundwater saturated (n=3).

### Metabolism prediction

Genes encoding for key enzymes in 23 metabolic functions were identified with 204 KOfam (Aramaki *et al.*, 2019) and custom Hidden Markov Models (Anantharaman *et al.*, 2016). Cutoffs for all custom models and for 10% of all KOfam models were tested and adjusted by searching for best matches of HMM hits against NCBI’s nr database using BLASTP. The search results and protein domains were compared with a reference protein sequence from Uniprot, and a stringent cutoff was selected to prevent false positives by setting it to be above the first false positive in the search results (Table S4). In case a genome was found to have >= 0.75 of all genes required to complete a function, and the genome was representing a dRep cluster, then the missing genes were sought for within the other genomes of the cluster using the same HMM model and thresholds. A metabolic function was considered present only if all required genes were present within a single genome (Table S4). The presence of *dsrD* gene was required in order to consider an organism, which has dsrAB genes, as a sulfide oxidizer. As some genomes lack *dsrD* due to genome incompleteness, concatenated *dsrAB* genes were aligned with a database of oxidizing and reducing *dsrAB* genes from Muller et al. (2015). The genes were aligned with MAFFT (Katoh and Standley, 2013) with default parameters, and a tree was inferred with Fasttree (Price *et al.*, 2010). Concatenated *dsrAB* genes that were located within the reductive clade were considered as potentially reductive *dsrAB*. The abundance of each function was determined by summing the average coverage of reads mapped to each organism that has all required genes to fulfill the function.

Due to structural similarities, the HMM which was designed to identify methane dehydrogenase (*mdh*) could also detect other alcohol dehydrogenases. The identity of the amino acid sequence was verified by placing the sequences along with a reference set into a phylogenetic tree as described by Diamond et. al. (Diamond *et al.*, 2019) and using the same reference set.

Hidden Markov Models for arsenic genes were custom made. The seed sequences for the HMMs were taken from Zhu et. al., (Zhu *et al.*, 2017) and references within. Seed sequences were added by searching for amino acid sequences of proteins with similar function and structure by searching NCBI’s nr database with BLASTp to increase the taxonomic diversity. For each gene, reference sequences from the same (or related) protein families were used to verify clustering of all seed sequences. These were taken from the NCBI Conserved Domain database. Once seed sequences were selected, HMMs were built with HMMER (Finn *et al.*, 2015). Score cutoffs were selected as reported above. To verify the results, sequences from the current study were aligned with the seed sequences and reference sequences using MAFFT (Katoh and Standley, 2013) and a FastTree was constructed to assess clustering.

### Taxonomy

The taxonomy of each genome was determined by comparing the concatenated sequences of 15 ribosomal proteins to a reference dataset (Hug *et al.*, 2013). The sequences of rpL2, rpL3, rpL4, rpL5 rpL6, rpL14, rpL15, rpL18, rpL22, rpS4, rpS8, rpS17 and rpS19 were identified with FetchMG v1.1. which is available as a standalone part of MOCAT (Kultima *et al.*, 2012). Additionally, rpS10 and rpL24 were searched for with TIGR01049 and TIGR01079 models for bacterial, or TIGR01046 and TIGR01080 models for archaeal bins, respectively (Haft *et al.*, 2001). The search was conducted with hmmsearch v3.1b2 from HMMER suite using NC cutoff implement within the model files (Finn *et al.*, 2015). The genes were concatenated and aligned to a consensus sequence generated from a 3078 sequences reference set and placed on a reference tree using a Neighbour-Joining (NJ) algorithm implemented in pplacer v1.1.alpha (Matsen *et al.*, 2010). Next, the 10 closest taxa to each query genome were identified and the genes of their 15 ribosomal proteins were extracted from the reference set. Each gene of the new, leaner reference set was aligned with the query gene using MAFFT (Katoh and Standley, 2013) and a Maximum-Likelihood (ML) tree was constructed on the CIPRES Science Gateway v3.3 (Miller *et al.*, 2010) with RAxML (Stamatakis, 2014) using the LG substitution model and bootstrapping, allowing the software to halt bootstrapping once it reached a consensus. The taxonomy of each query genome was inferred based on the topology of the tree and its closest reference genome.

**CH_4_ concentrations** were quantified on a GC-2014 Shimadzu gas chromatograph (Shimadzu Corporation, Kyoto, Japan). Using a gas-tight syringe, 4.5 mL of sample aliquot was injected into a 1 mL stainless steel loop mounted on a 10-port valve (Valco). CH_4_ was separated on a packed HayeSep-D packed column (4 m × 1/8 inch). CH_4_ was quantified with a flame ionization detector (FID). For water samples, after headspace concentration measurements, methane dissolved in the water concentrations were calculated using Henry’s Law constant. Analytical precision based on repeated standard analyses was ~ 3% of the reported concentrations.

**Carbon and hydrogen isotope ratios of CH_4_** were measured separately using a gas chromatograph and an isotope ratio mass spectrometer interfaced with a pyrolysis (GC-P-IRMS) and a combustion reactor (GC-C-IRMS, Thermo Fisher Scientific, Bremen, Germany). CH_4_ was separated chromatographically on an HP-molesieve fused silica capillary column (30 m × 0.320 mm). For hydrogen isotopes, after GC separation, CH_4_ was pyrolyzed in a capillary carbon-coated ceramic tube at 1450 °C and the hydrogen isotope ratios were measured in the IRMS. For carbon isotopes, after GC separation, the CH_4_ was combusted to CO_2_ at 1030 °C in a capillary ceramic tube loaded with Ni, Cu, and Pt wires and the carbon isotope ratio was acquired in the IRMS. Results are reported in the standard δ notation as the per mil deviation (‰) relative to Vienna Pee Dee Belemnite (VPDB) for carbon and relative to Vienna Standard Mean Ocean Water (VSMOW) for hydrogen. Repeated injections of CH_4_ yield values of −39.46 ± 0.33‰ (1σ; n=11) and −167.01 ± 3.10‰ (1σ; n=14) for δ^2^H.

### Statistical analysis

All analysis was done R v3.4.3 (R Development Core Team, 2012) and Rstudio v1.1.423 (Rstudio Team, 2015). Correlation between genomic bins and sampling zones was done with “multipatt” command and r.g. function in the Indicspecies package v1.7.6 (Caceres and Legendre, 2009). Indicator function was tested with the IndVal.g function. In both cases, 9999 permutations were conducted and the p-value was corrected for multiple testing with Benjamini-Hochberg procedure. Maps were retrieved from Google maps database using Google Earth v7.3.2.

## Availability of data and material

- Raw reads are available through the NCBI Short Reads Archive. Accession number for each sample is provided in Table s1.
- Sequences of the 484 Metagenomic Assembled Genomes are available at: https://ggkbase.berkeley.edu/PLM2016-NC-80p/organisms
- HMMs used in the current study (ref to the updated list, and list of ko and thresholds for KOfam) are available at https://figshare.com/account/home#/projects/66182
- Phylogenetic tree of 15RP https://figshare.com/account/home#/projects/66182
- Presence/absence of genes in each genome and the rules by which presence/absence of fucuntions were determined. https://figshare.com/account/home#/projects/66182

## Authors’ contributions

A.L designed research, performed research, analyzed data, and wrote the paper. P.B.M.C conducted fieldwork and assisted in constructing the metabolic pathway analysis. R.K created and tested Hidden Markov Models for arsenic metabolism related genes. M.B. contributed to methane data acquisition and in writing the paper. J.W assisted in designing and conducting fieldwork. T.K.T assisted in designing and conducting fieldwork. K.H.W took part in the research design and assisted in fieldwork. S.S.H took part in the research design. J.F.B supervised the study and mentored the first author.

## Supplement material

**Figure S1.**
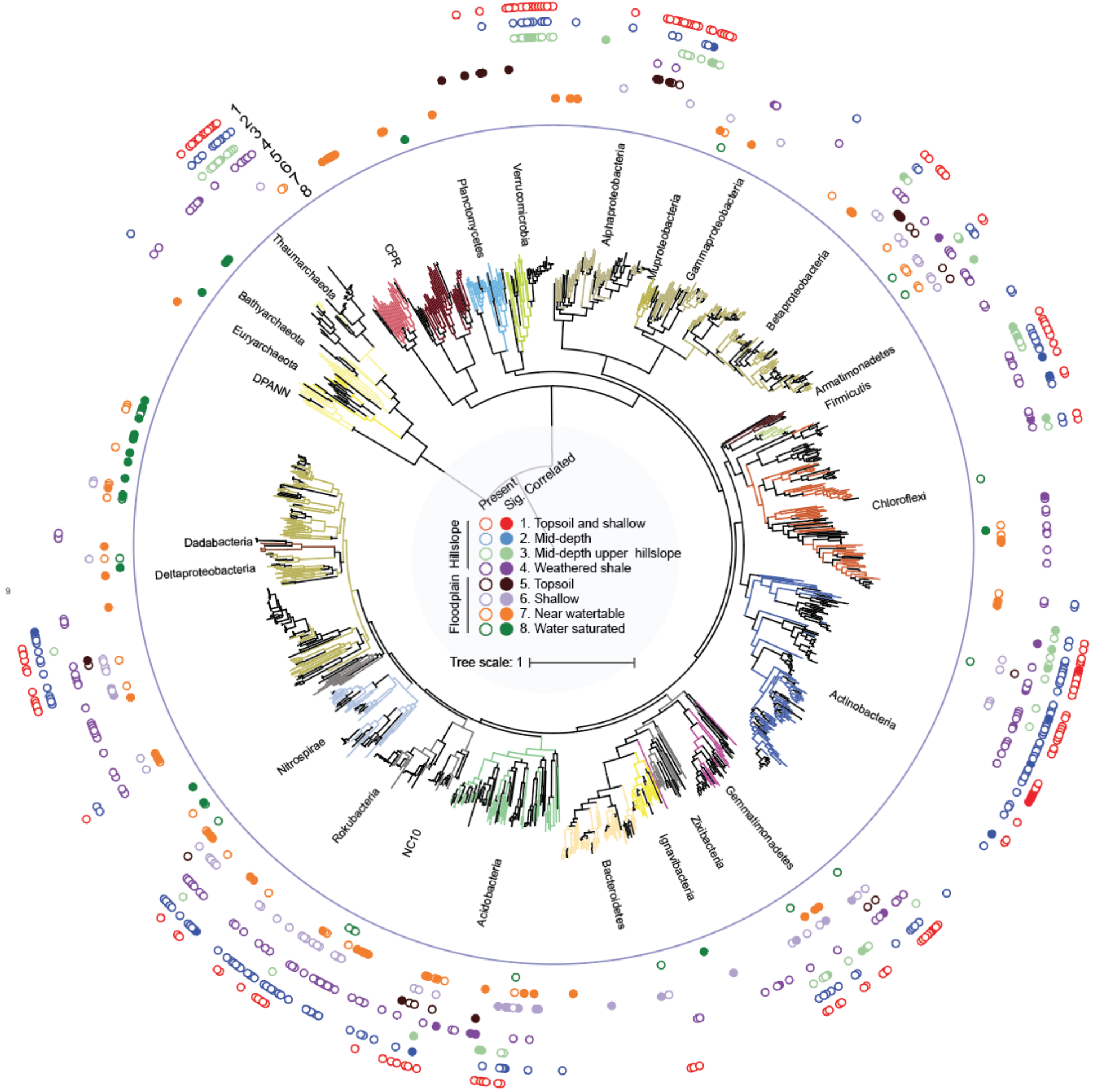
Correlation of microbial species with ecosystem compartment along hillslope to Riparian-zone transect. A maximum-likelihood tree of 16 Ribosomal Proteins of 484 representative genomes (black lines) and reference genomes (colored lines). The presence of a genome at each location is denoted by an empty circle. A species found to be correlated with a specific sample is marked with a filled circle (IndVal analysis, p-value < 0.05, 9999 permutations, Benjamini-Hochberg corrected).

**Figure S2.**
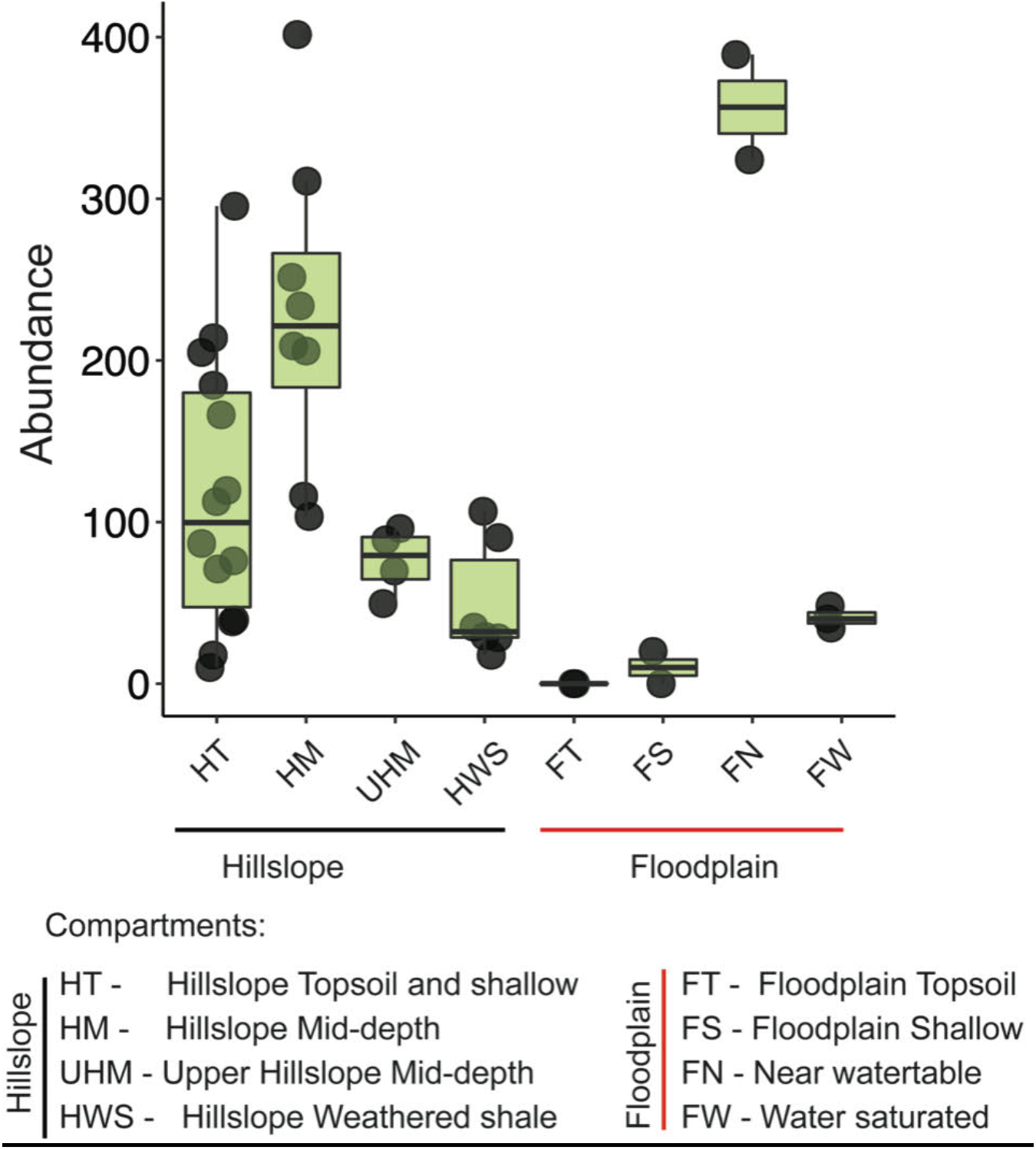
Abundance of organisms with arsenic methylation related genes. Each dot represents one sample. Samples are clustered spatially by compartment

**Figure S3:**
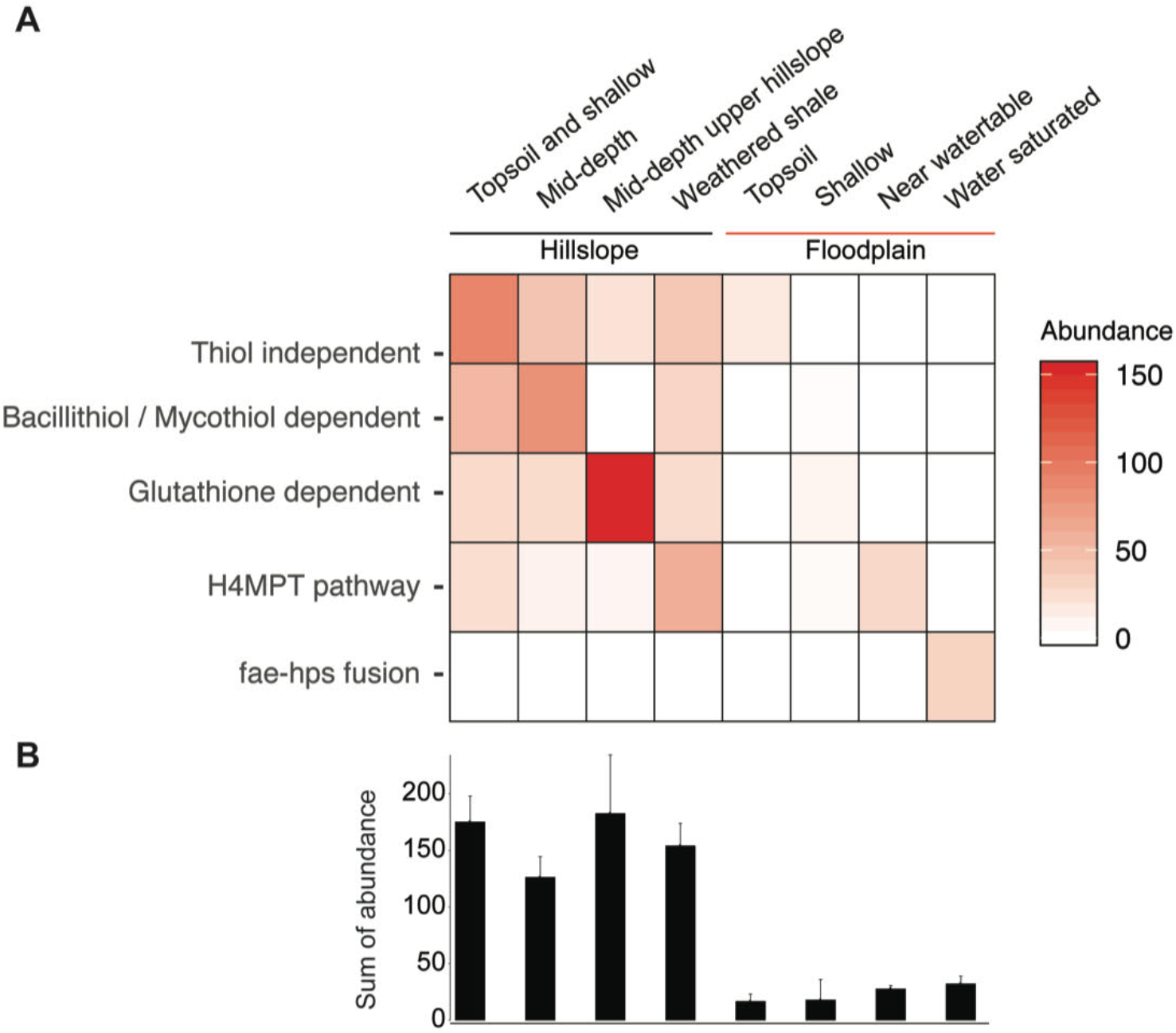
Abundance of organisms with the different formaldehyde oxidation pathways detected across the hillslope to riparian zone transect. **A. The diversity and abundance of genomes with formaldehyde oxidation pathways across the study sites.**The abundance depicted in each box is the sum abundance of all genomes that have the function in a specific site. B. Sum abundance of genomes that have the capacity for **formaldehyde oxidation at each site**.

**Figure S4:**
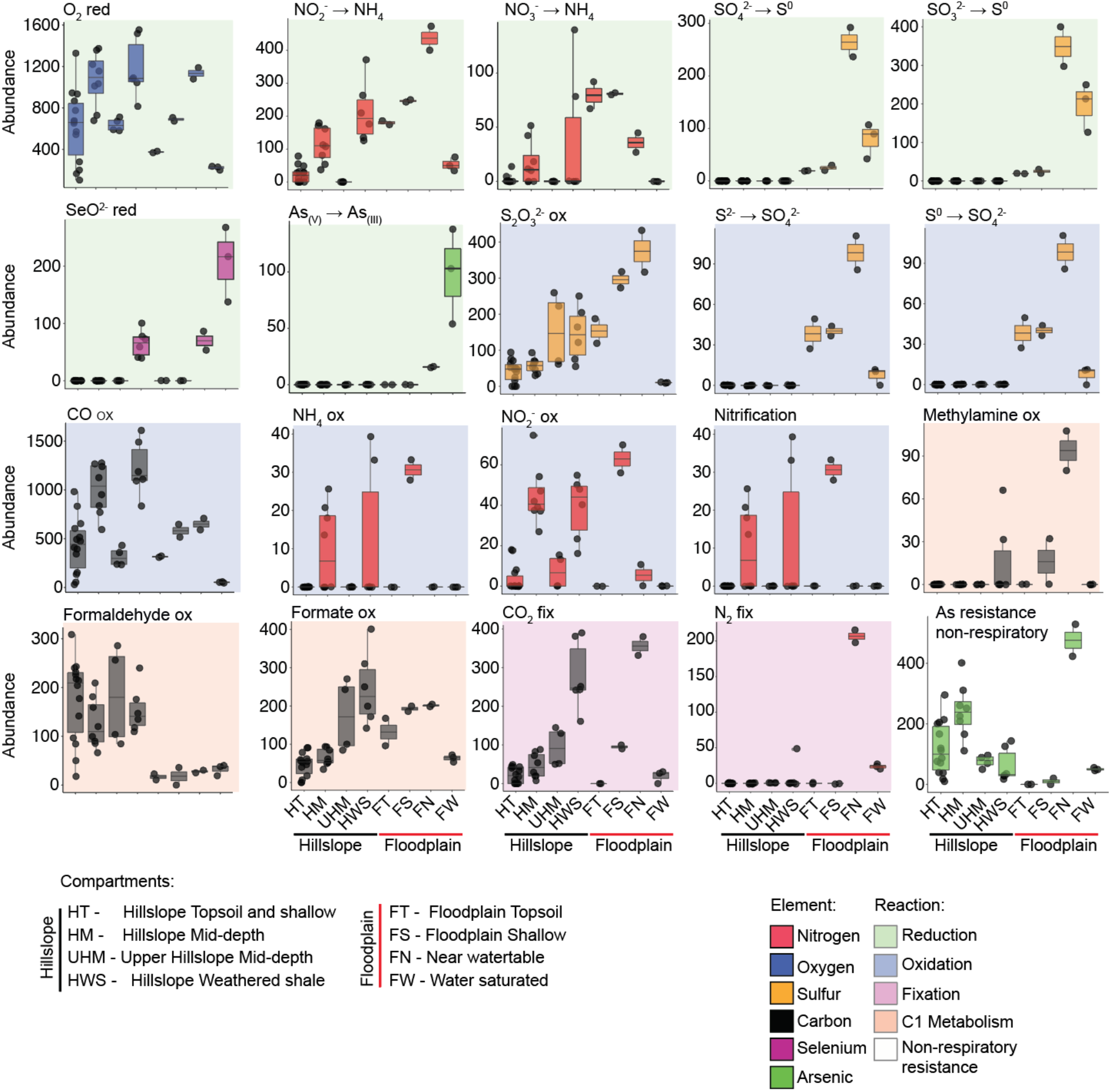
The abundance of metabolic functions. Middle bar of each box plot represents the mean abundance of organisms at this location. The color of each bar represents the element which is directly involved in the metabolic pathway. The type of reaction (i.e., C1 metabolism, reduction, oxidation, fixation or non-respiratory resistance) is denoted by the color of background behind the bars.

**Figure S5:**
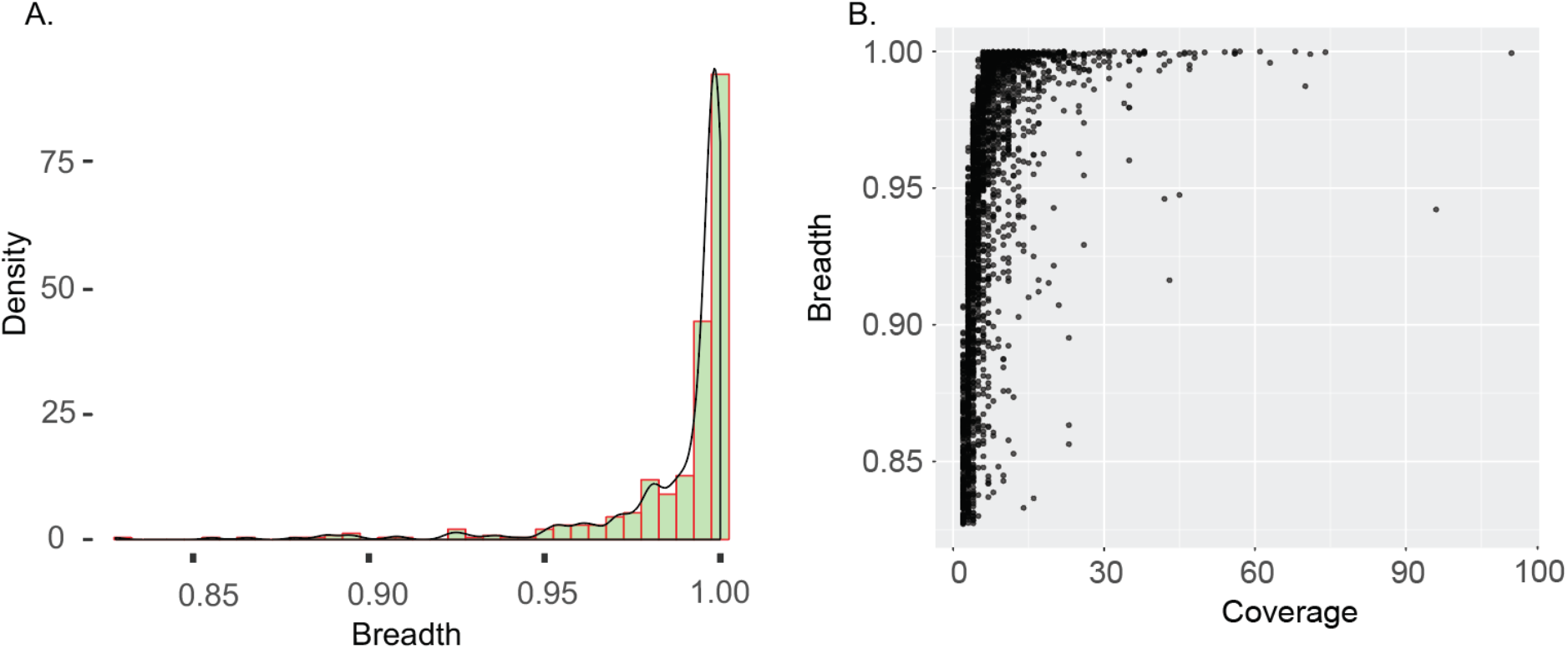
Minimal breadth required in order to consider a genome present in a sample. A. The breadth of genomes as calculated by mapping reads of the sample from which the genome was binned. An 82.8% breadth was found to be the minimum required to obtain a genome from its original sample. B. The breadth and coverage of all genomes passing the 82.8% breadth cutoff are shown. Each dot represents one genome.

